# ω-Phonetoxins inhibit voltage-gated calcium Ca_V_2.2 ion channel splice isoforms of dorsal root ganglia

**DOI:** 10.1101/2023.08.29.555337

**Authors:** Célio Castro-Junior, Marcus Vinícius Gomez, Marcia Helena Borges, Diane Lipscombe, Arturo Andrade

## Abstract

Cell-specific alternative splicing of *Cacna1b* pre-mRNA generates functionally distinct voltage-gated Ca_V_2.2 channels. Ca_V_2.2 channels mediate the release of glutamate from nociceptor termini in the dorsal horn spinal cord and they are implicated in chronic pain. One alternatively spliced exon in *Cacna1b*, e37a, is highly expressed in dorsal root ganglia, relative to other regions of the nervous system, and it is particularly important in inflammatory hyperalgesia. Here we studied the effects of two ω-phonetoxins, PnTx3-4 and Phα1β, derived from the spider *Phoneutria nigriventer* on Ca_V_2.2 channel isoforms of dorsal root ganglia (Ca_V_2.2 e37a and Ca_V_2.2 e37b). Both PnTx3-4 and Phα1β are known to have analgesic effects in rodent models of pain and to inhibit Ca_V_2.2 channels. Ca_V_2.2 e37a and Ca_V_2.2 e37b isoforms expressed in a mammalian cell line were inhibited by PnTx3-4 and Phα1β with similar potency and with similar timecourse, although Ca_V_2.2 e37a currents were slightly, but consistently more sensitive to toxin inhibition compared to Ca_V_2.2 e37b. The inhibitory effects of PnTx3-4 and Phα1β on Ca_V_2.2-e37a and Ca_V_2.2-e37b channels were voltage-dependent, and both occlude the inhibitory effects of ω-conotoxin GVIA, consistent with a common site of action. The potency of PnTx3-4 and Phα1β on both major splice isoforms in dorsal root ganglia constribute to understanding the analgesic actions of these ω-phonetoxins.

## 1. Introduction

Presynaptic voltage-gated calcium Ca_V_2.2 ion channels mediate calcium entry that triggers transmitter release at many synapses. At presynaptic termini in spinal cord dorsal horn, Ca_V_2.2 channels triggers the release of glutamate and substance P {Bertolino and Llinás, 1992, #38408; Holz et al., 1988, #2015; Malmberg and Yaksh, 1994, #299456; Scroggs and Fox, 1992, #181759; Scroggs and Fox, 1992, #167853; Zamponi et al., 2009, #65832}. Mice lacking Ca_V_2.2 channels have impaired responses to noxious stimuli {Kim et al., 2001, #65057; Hatakeyama et al., 2001, #108707; Saegusa et al., 2001, #225859; DuBreuil et al., 2021, #163505} and, in the clinic, the Ca_V_2.2 channel peptide toxin ω-Conotoxin MVIIA (ω-CTX MVIIA) ameliorates refractory pain in patients with cancer and AIDS {Staats et al., 2004, #276689}.

Ca_V_2.2 is encoded by *Cacna1b*, a multi-exon gene with several sites of alternative splicing (e18a, e24a, e31a and 37a/37b) {Lipscombe et al., 2013, #3999; Lipscombe et al., 2013, #286100}. Native Ca_V_2.2 channel currents represent the collective activation of a subset of Ca_V_2.2 channel splice isoforms, the composition of which depends on cell-type, development and localization {Lipscombe and Lopez Soto, 2019, #110892; López Soto and Lipscombe, 2020, #223520}. In nociceptors two major forms of Ca_V_2.2 dominate, which are distinguished based on the presence of either e37a or e37b encoding sequence. Ca_V_2.2-e37a channels have highly restricted expression profile in *transient receptor potential vanilloid 1 (Trpv1)*-lineage nociceptors {Bell et al., 2004, #275808; López Soto and Lipscombe, 2020, #223520} and they are more sensitive to inhibition by G protein coupled receptors including μ-opioid receptors {Andrade et al., 2010, #78806; Raingo et al., 2007, #119323}. The presence of Ca_V_2.2-e37a channels in nociceptors underlies the unique properties of native Ca_V_2.2 current in these neurons {Altier et al., 2007, #229448; Andrade et al., 2010, #78806; Bell et al., 2004, #275808; Castiglioni et al., 2006, #182531; Jiang et al., 2013, #131588; López Soto and Lipscombe, 2020, #223520; Macabuag and Dolphin, 2015, #207013}.

Ca_V_2.2-e37a channel isoforms are implicated in inflammatory thermal hyperalgesia in mice {Altier et al., 2007, #229448}, suggesting that Ca_V_2.2-e37a might be a particularly important therapeutic target to treat certain forms of chronic pain. In addition, in animal models of neuropathic pain, *Cacna1b* e37a mRNA splice variants are reduced in *Trpv1-lineage* nociceptors {López Soto and Lipscombe, 2020, #223520}, likely contributing to the reduced analgesic actions of morphine in chronic pain {Andrade et al., 2010, #78806; Jiang et al., 2013, #131588}.

We tested ω-phonetoxins from the venom of the spider *P. nigriventer* {Gomez et al., 2002, #1230; Peigneur et al., 2018, #155522; Peigneur et al., 2018, #59374} for their ability to inhibit Ca_V_2.2 channel splice isoforms. Of 40 peptide toxins derived from *P. nigriventer*, PnTx3-4 (previously known as ω-ctenitoxin-Pn3a) and Phα1β (previously named PnTx3-6 or ω-ctenitoxin-Pn4a) block high voltage activated calcium channels including Ca_V_2.2 {Cassola et al., 1998, #125214; Vieira et al., 2005, #270589}, and these toxins are analgesic in animal models of chronic pain {da Silva et al., 2015, #70393; de Souza et al., 2013, #23934; Diniz et al., 2014, #91186; Rigo et al., 2013, #61858; Rigo et al., 2013, #154046; Souza et al., 2008, #233135; Tonello et al., 2014, #159900}. PnTx3-4 inhibits all three Ca_V_2 channels, Ca_V_2.1, Ca_V_2.2, and Ca_V_2.3, with small differences in potency {Dos Santos et al., 2002, #127690}, whereas Phα1β inhibits Ca_V_2.2 with greater potency as compared to Ca_V_2.3, Ca_V_2.1 and Ca_V_1.2 channels {Vieira et al., 2005, #270589}.

We find that PnTx3-4 and Phα1β inhibit Ca_V_2.2-e37a and Ca_V_2.2-e37b channel isoforms with similar efficacy although Ca_V_2.2-e37a currents activated at relatively negative voltages, are slightly more sensitive to inhibiton by a submaximal concentration of PnTx3-4 compared to Ca_V_2.2-e37b.

## 2. Results

ω-CTX GVIA is a highly specific inhibitor of Ca_V_2.2, as previously reported {Jiang et al., 2013, #131588}, we show that ω-CTX GVIA inhibits both Ca_V_2.2-e37a and Ca_V_2.2-e37b splice isoform currents expressed in tsA201 cells equally well, and at 1 μM ω-CTX GVIA inhibits 100 % of Ca_V_2.2 currents (**Fig. 1A-C**). We next compared the inhibitory actions of PnTx3-4 and Phα1β toxins on Ca_V_2.2-e37a and Ca_V_2.2-e37b currents. At 500 nM, PnTx3-4 inhibited Ca_V_2.2-e37a and Ca_V_2.2-e37b currents by 76.8 ± 1.7 % (n = 5) and 66.6 ± 6.2 % (n = 4), respectively (**Fig. 1D-F**; p = 0.123, Student’s t-test) and 500 nM Phα1β inhibited currents by 75.2 ± 3.6 % (n = 7) and 64.7 ± 5.2 % (n = 7), respectively (Fig. **1G-I**; p = 0.124, Student’s t-test). The inhibitory actions of PnTx3-4 and Phα1β on Ca_V_2.2 currents were long lasting and effectively irreversible after 100 s of continuous wash (**Fig. 1E** and **1H**).

**Figure 1:**
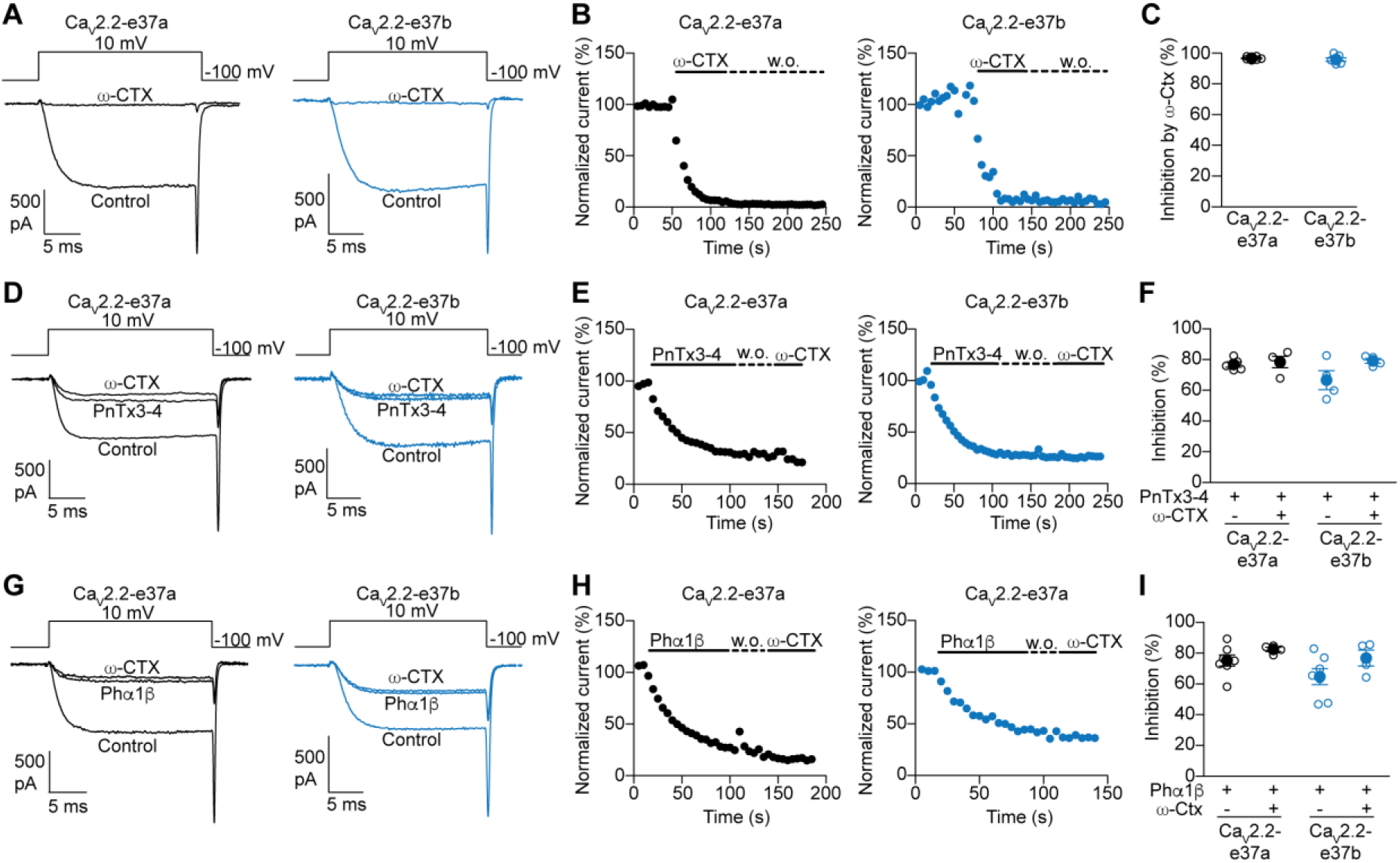
ω-Phonetoxins, PnTx3-4 and Phα1β, inhibit Ca_V_2.2-37a and Ca_V_2.2-37b splice isoforms and occlude the actions of ω-conotoxin GVIA. **A-C**) Action of 1 μM ω-conotoxin GVIA (ω-CTX GVIA) on Ca_V_2.2-37a and Ca_V_2.2-37b currents. Example Ca_V_2.2 current recordings (A), time course of inhibition (B), and percent inhibition (C), for 1 μM ω-CTX GVIA on Ca_V_2.2-37a and Ca_V_2.2-37b splice isoforms. In C, individual data points (open symbols) and average values (closed symbols) summarizing percent inhibition at steady state block are shown. **D-I**) Action of 500 nM PnTx3-4 (D-F) or Phα1β (G-I) and 1 μM ω-CTX on Ca_V_2.2-37a and Ca_V_2.2-37b currents. PnTx3-4 or Phα1β was applied for ∼60 s, then the cells were washed with control solution for 100 s, followed by application of μM ω-CTX for another 40 s. Example Ca_V_2.2 current recordings (D, G), time course of inhibition (E, H), and percent inhibition of 7 cells (F, I), for 500 nM PnTx3-4 or Phα1β and 1 μM ω-CTX block of Ca_V_2.2-37a and Ca_V_2.2-37b splice isoforms. In F, I, individual data points (open symbols) and average values (closed symbols) summarizing percent inhibition at steady state block are shown. Toxin application is indicated with bold horizontal lines and wash out (w.o.) with dashed horizontal lines. All Ca_V_2.2 currents were evoked by step depolarizations from -100 mV to 10 mV applied at 5 sec intervals in tsA201 cells.

ω-CTX GVIA and ω-CTX MVIIC can displace ω-phonetoxins, including ω-phonetoxin IIA (ω-PtxIIA), from synaptosome preparations containing Ca_V_2.1, Ca_V_2.2 and Ca_V_2.3 channels {Dos Santos et al., 2002, #29669} and the high affinity binding site for ω-PtxIIA appear to partially overlap with ω-CTX GVIA {Dos Santos et al., 2002, #127690}. Consistent with these reports we show that PnTx3-4 occludes the actions ω-CTX GVIA. When applied following 500 nM PnTx3-4, which inhibited 76.8 ± 1.7 % of Ca_V_2.2-e37a current (**Fig. 1F**) and 66.6 ± 6.2 % of Ca_V_2.2-e37b current (**Fig. 1F**), 1 μM ω-CTX GVIA had limited effect at a concentration which normally inhibits 100% of Ca_V_2.2 currents (**Fig. 1C**) {Jiang et al., 2013, #131588} (Ca_V_2.2-e37a: p = 0.606, n = 4; Ca_V_2.2-e37b: p = 0.155, Student’s paired t-test, n = 4) (**Fig. 1D-F**). 1 μM ω-CTX GVIA was similarly ineffective when applied after Phα1β. At 500 nM, of Phα1β inhibited 75.1 ± 3.6 % of Ca_V_2.2-e37a current compared to 82.4 ± 1.4 % in the presence of 1 μM ω-CTX GVIA (p = 0.31, Student’s paired t-test, n = 4). At 500 nM, Phα1β inhibited 64.7 ± 5.2 % of Ca_V_2.2-e37b current compared to 76.8 ± 5.1 % in the presence of 1 μM ω-CTX GVIA (p = 0.1140, Student’s paired t-test, n = 4) (**Fig. 1G-I**).

At increasing concentrations, PnTx3-4 (**Fig. 2A–C**) and Phα1β (**Fig. 2D–F**) inhibited Ca_V_2.2 splice isoform currents incrementally: the IC_50_ for PnTx3-4 inhibition of Ca_V_2.2-e37a currents was 59 nM (95% CI: 16 – 213 nM) and for Ca_V_2.2-e37b was 107 nM (95% CI: 21 – 535 nM) (p = 0.44, Students t-test) (**Fig. 2C)**; and the IC_50_ for Phα1β inhibition of Ca_V_2.2-e37a was 211 nM (95% CI: 29 – 1513 nM) and for Ca_V_2.2-e37b was 142 nM (95% CI: 27 – 230 nM) (p = 0.44, Student’s t-test) (**Fig. 2F)**.

**Figure 2:**
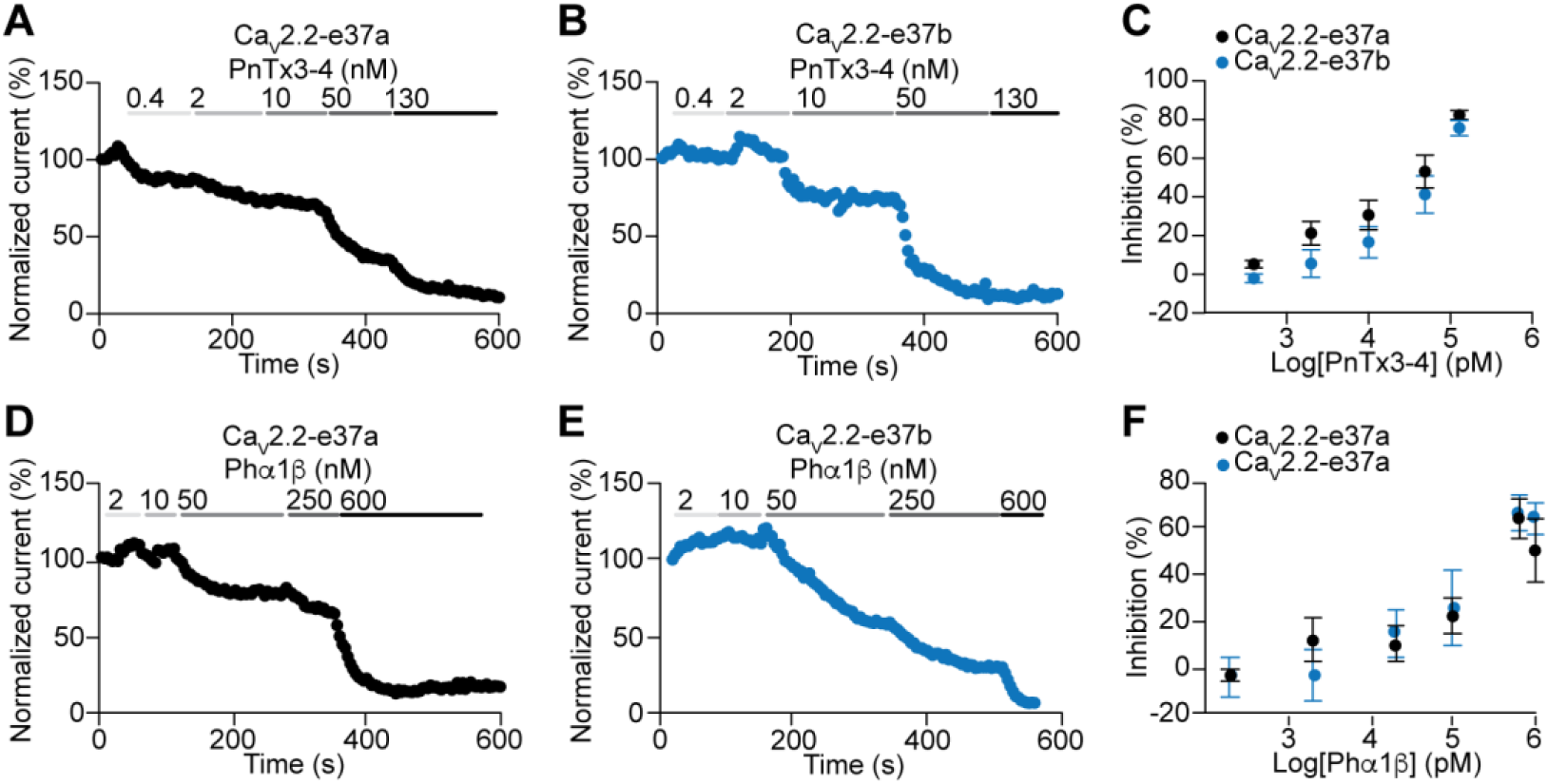
Dose-response curves for PnTx3-4 and Phα1β inhibition of Ca_V_2.2-37a and Ca_V_2.2-37b currents. Effect of increasing concnetrations of PnTx3-4 (**A-C**) and Phα1β (**D-F**) on Ca_V_2.2-37a (left, black lines/symbols) and Ca_V_2.2-37b (right, blue lines/symbols) currents recorded in tsA201 cells. Example time courses (A, B, D, E) and average inhibition (C, F) are shown. C, F, Average percent inhibition of at least 5 cells per point are showing together with standard errors .

The actions of peptide toxins and other inhibitors of voltage gated ion channels are reported to depend on the state of the channels – e.g. closed, open or inactivated {Catterall et al., 2007, #243689; Feng et al., 2003, #162567; McDonough et al., 1997, #46854; Peigneur et al., 2018, #155522}. Both PnTx3-4 and Phα1β exhibit some degree of voltage-dependence to their inhibitory actions on Ca_V_2.2-e37a and Ca_V_2.2-e37b splice isoforms (**Fig. 3C** and **3F**), but this was more pronounced with Phα1β. At negative membrane voltages (-20 mV) currents were only ∼40% of peak amplitude in the presence of Phα1β compared to ∼ 70% of peak amplitude at 0 mV. In addition, Ca_V_2.2-e37a currents were more sensitive to inhibition by Phα1β compared to Ca_V_2.2-e37b currents particularly when activated by voltages between -20 and -5 mV (ANOVA repeated measures, F_(1,12)_ = 136.22, p < 0.0001, **Fig. 3F**), but this difference in toxin action was substantially reduced on currents evoked by stronger depolarizations to 0 and 40 mV (ANOVA repeated measures, F_(1,12)_ = 2.572, p = 0.135, **Fig. 3F**). Our results suggest that the effects of PnTx3-4 and Phα1β on Ca_V_2.2-e37a and Ca_V_2.2-e37b splice isoforms are voltage-dependent and that Ca_V_2.2-e37a currents currents evoked at more negative membrane voltages ∼ -20 mV are slightly more sensitive to Phα1β as compared to Ca_V_2.2-e37b (**Fig. 3F**)

**Figure 3:**
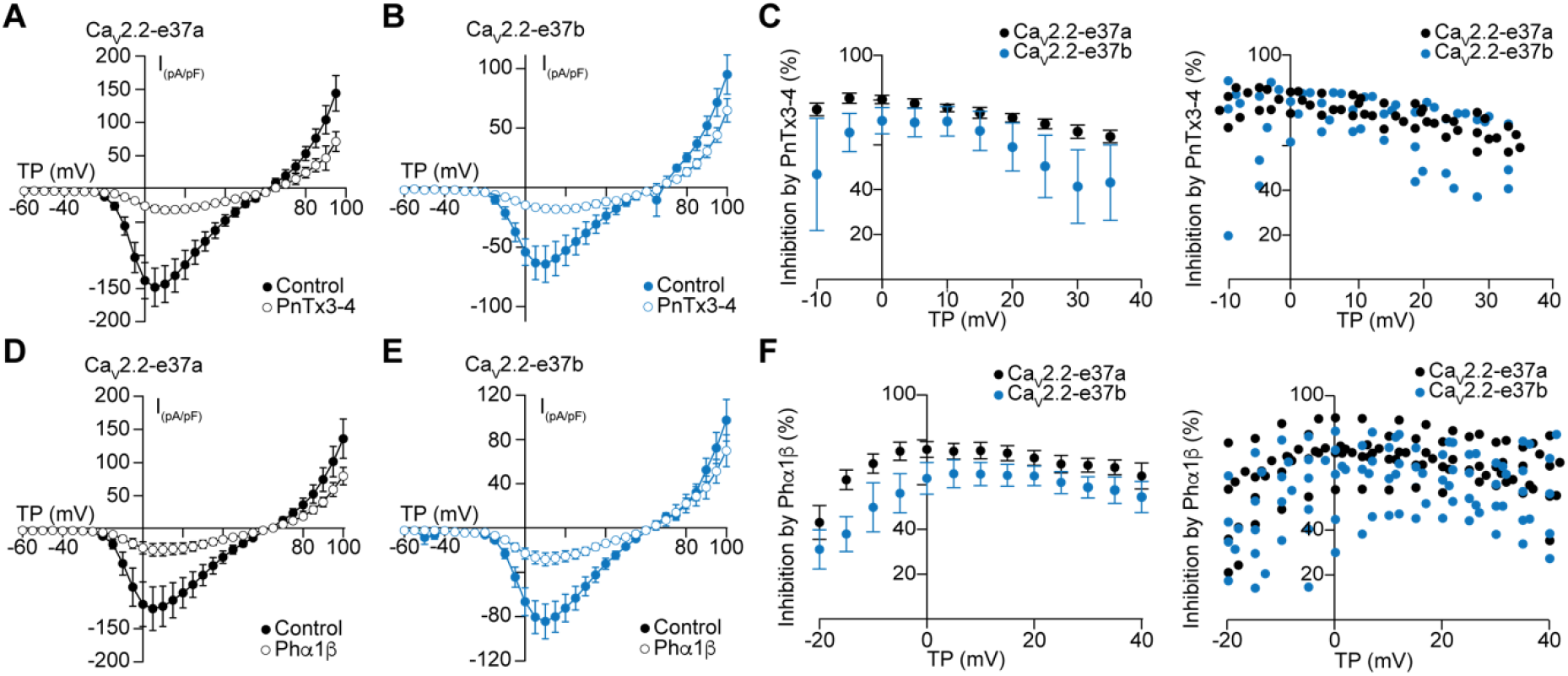
Voltage-depedent inhibition of Ca_V_2.2-e37a and Ca_V_2.2-e37b isoforms by ω-phonetoxins. Voltage dependent actions of 500 nM PnTx3-4 (A-C) and 500 nM Phα1β (D-F) on Ca_V_2.2-e37a and Ca_V_2.2-e37b currents evoked by voltage steps between -60 mV and +100 mV in 5 mV increments, from a holding potential of -100 mV. Average current voltage relationships (A, B, D, E), average inhibtion (left) and inhibition of individual recordings (right) at different test potentials (C, F) are shown. Average data shown are mean ± SE. ± SE.

Finally, we compared the time course of toxin inhibition of Ca_V_2.2-e37a and Ca_V_2.2-e37b currents using submaximal concentrations of PnTx3-4 and Phα1β (140 nM). Solution exchange occurred within 1-2 seconds and steady state inhibition of currents was achieved within 50-80 sec. The time courses of PnTx3-4 and Phα1β inhibition of Ca_V_2.2-e37a and Ca_V_2.2-e37b currents were overall similar (PnTx3-4: τ_on_ median and IQR: Ca_V_2.2-e37a = 50.77 s, 40.33 s, n = 16; Ca_V_2.2-e37b = 69.18 s, 102.52 s, n = 18. Wilcoxon Rank Sum test, p = 0.07, **Fig. 4A-C;** Phα1β: τ_on_ mean ± s.e.m: Ca_V_2.2-e37a = 69.6 ± 18.7 s, n = 7; e37b-*Cacna1b* = 94.3 ± 23.1 s, n = 7. Student’s t-test, p = 0.422. **Fig. 4E-G**) but, as we observed in other experiments toxin inhibition of Ca_V_2.2-e37a currents was slightly but consistently greater as compared to Ca_V_2.2-e37b (PnTx3-4 % inhibition and IQR: Ca_V_2.2-e37a = 78.3, 10.03, n = 18; Ca_V_2.2-e37b = 74.33, 10.6, n = 18, Wilcoxon Rank Sum test, p = 0.01, **Fig 4D;** Phα1β % inhibition ± s.e.m: Ca_V_2.2-e37a = 75.2 ± 3.6, n = 7; Ca_V_2.2-e37b = 64.7 ± 5.2, n = 7. Student’s t-test, p = 0.1236. **Fig. 4H**).

**Figure 4:**
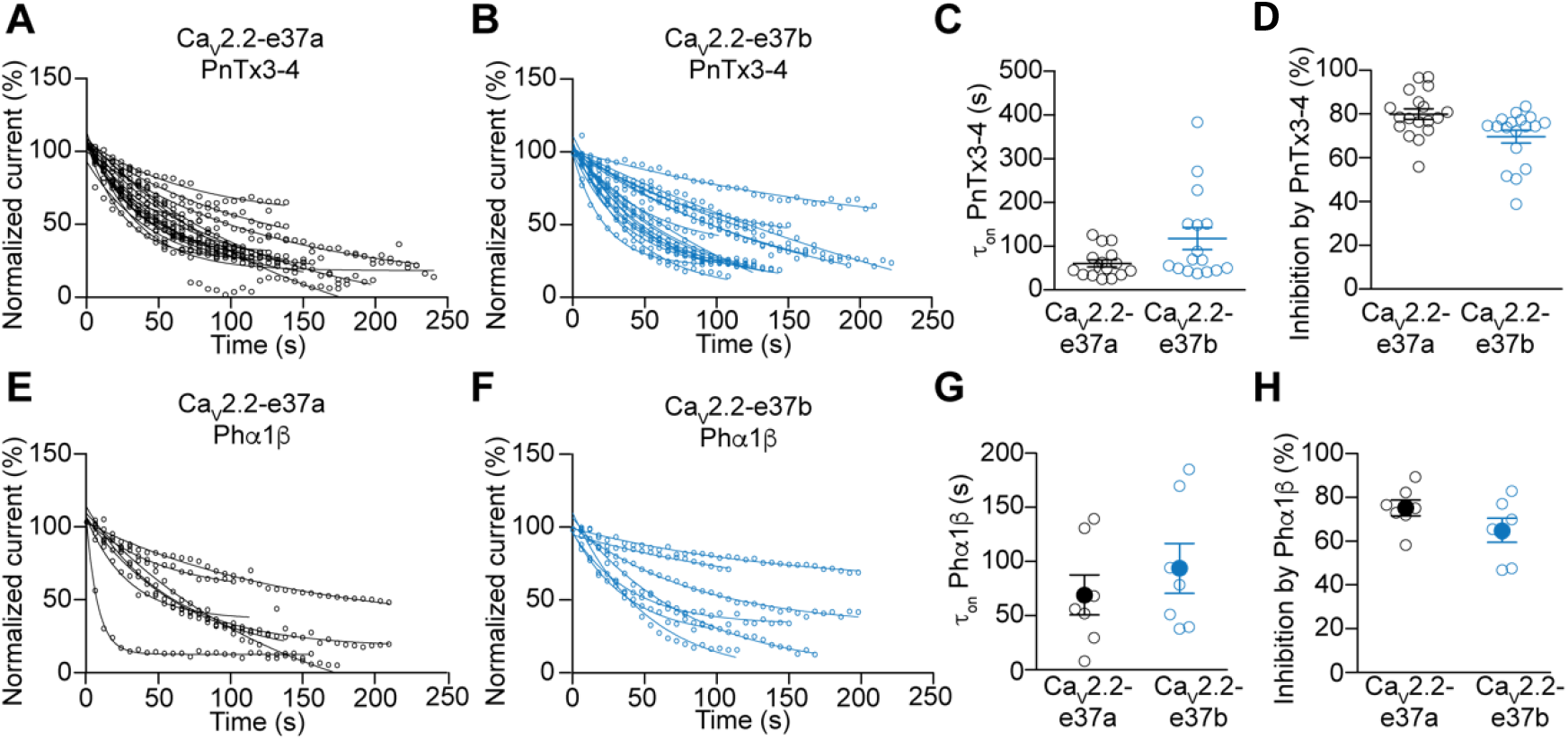
ω-Phonetoxins inhibit Ca_V_2.2-e37a and Ca_V_2.2-e37b currents with similar time course. **A-H)**, Summary of the time course of inhibition of Ca_V_2.2-e37a and Ca_V_2.2-e37b currents by submaximal concentration (140 nM) of PnTx3-4 (A-D) and Phα1β (E–H). Time course of toxin inhibition of individual recordings (A, E), τ_on_ values from single exponential fits to individual data (D, G), and percental inhibition are shown. C, D, data from each cell (open symbols) are shown together with boxplots representing the interquartile range (IQR), median, and 95% confidence intervals. G, H, data from each cell (open symbolts) are shown together mean ± SE.

## 3. Discussion

We report the effects of two toxins, PnTx3-4 and Phα1β, obtained from the venom of the spider *P. nigriventer* on Ca_V_2.2-e37a and Ca_V_2.2-e37b currents expressed in a mammalian cell line. We find PnTx3-4 and Phα1β inhibit Ca_V_2.2-e37a and Ca_V_2.2-e37b with similar efficacy although Phα1β is slightly more effective on Ca_V_2.2-e37a currents evoked at the foot of the current-voltage relationship (∼ -20 mV) compared to Ca_V_2.2-e37a. Ca_V_2.2-e37a and Ca_V_2.2-e37b splice isoforms are both expressed in sensory neurons of dorsal root ganglia and, Ca_V_2.2-e37a is specifically enriched in *Trpv1*-lineage neurons. The ability of spider *P. nigriventer* toxins to inhibit both Ca_V_2.2-e37a and Ca_V_2.2-e37b currents in dorsal root ganglia contributes to their analgesic properties.

PnTx3-4 and Phα1β inhibition of Ca_V_2.2-e37a and Ca_V_2.2-e37b currents was incomplete (maxium 70-80% inhibition at 500 nM), in constrast to the conus toxin ω-CTX GVIA which inhibits close to 100% of Ca_V_2.2 currents (**Fig. 1**) {Jiang et al., 2013, #131588}. Our findings in this respect are consistent with analyses of Phα1β on Ca_V_2.2 currents reported previously by Vieira and colleagues {Vieira et al., 2005, #270589} although PnTx3-4 was found to have greater potency against Ca_V_2.2 currents expressed in baby hamster kidney (BHK) cells {Dos Santos et al., 2002, #127690}. Similar to dos Santos and colleagues {Dos Santos et al., 2002, #127690}, we find that PnTx3-4 and Phα1β occlude the inhibitory actions of ω-CTX GVIA on calcium currents indicative of a common binding site. ω-CTX GVIA is widely used as a highly specific, and potent inhibitor of Ca_V_2.2. ω-CTX GVIA binds to the external loop between the membrane-spanning segment S5 and the pore-lining segment H5 in domain III to inhibit the ion channel pore {Boland et al., 1994, #11356; Ellinor et al., 1994, #255896}.

We now know that there is substantial diversity in the specific properties of major ion channel currents, including Ca_V_2.2, across cell types and these differences affect the sensitivity of Ca_V_2.2 channels to neurotransmitters and drugs. Specifically, Ca_V_2.2-e37a splice isoforms are more sensitive to inhibiton by G_i/o_ protein coupled receptor activation, explaining the high sensitivity of Ca_V_2.2 currents in *Trpv1*-nociceptors, which express relatively high levels of Ca_V_2.2-e37a splice isoforms, to morphine and other μ-opioid receptor agonists {Andrade et al., 2010, #78806; Bell et al., 2004, #275808; López Soto and Lipscombe, 2020, #223520; Raingo et al., 2007, #119323}. The inhibitory actions of ω-phonetoxins on Ca_V_2.2-e37a and Ca_V_2.2-e37b splice isoforms were overall similar, although we did observe small, but consistently higher potency against Ca_V_2.2 e37a compared to Ca_V_2.2 e37b. ω-Phonetoxins likely bind close to the ion pore, a region that is not different between Ca_V_2.2-e37a and Ca_V_2.2-e37b isoforms. However, Ca_V_2.2-e37a and Ca_V_2.2-e37b isoforms which differ in the composition of their C-termini, open at slightly different membrane voltages; Ca_V_2.2-e37a channels open at membrane voltages ∼ 10 mV more negative compared to Ca_V_2.2-e37b {Castiglioni et al., 2006, #182531}. These biophysical properties might explain the slightly greater potency of ω-Phonetoxins on Ca_V_2.2 e37a compared to Ca_V_2.2 e37b channel currents at relatively negative voltages, where the isoforms have the largest difference in activation thresholds.

Our studies shed light on the analgesic actions of ω-Phonetoxins by demonstrating that these toxins inbibit both major Ca_V_2.2 splice isoforms that are expressed in nociceptors. We observed only small differences in toxin action on Ca_V_2.2-e37a and Ca_V_2.2-e37b isoforms which correlate with small differences in voltage activation thresholds.

## 5. Methods

### 5.1 Toxins

Native PnTx3-4 and Phα1β were purified using a combination of gel filtration, reverse-phase fast protein liquid chromatography (GE, MN, USA) and high-performance liquid chromatography (Shimadzu, Tokyo, Japan) as described previously {Cordeiro et al., 1993, #245514; Scroggs and Fox, 1992, #181759}. PhTx3-4: SCINVGDFCDGKKDCCQCDRDNAFCSCSVIFGYKTNCRCEVGTTATS YGICNAKHKCGRQTTCTKPCLSKRCRRNHG (8419 Da, NCBI P81790). Phα1β: ACIPRGEICTDDCECC GCDNQCYCPPGSSLGIFKCSCAHANKYFCNRKKEKCKKA (6045 Da, NCBI AAB25575). The snail toxin ω-CTX GVIA was obtained from Alomone (Jerusalem, Israel).

### 5.2 Transient expression

tsA201 cells were maintained at 37°C with 5% CO_2_ in 90% Dulbecco’s modified Eagle’s medium (DMEM), supplemented with 10% fetal bovine serum (FBS). Variation in expression was minimized by selecting cells at 70% confluence for transfection, controlling cDNA concentrations, and harvesting all cells on the day of recording (24 h after transfection) and maintaining them in DMEM at 4°C until needed. We expressed mammalian cDNAs encoding calcium-channel e37a-*Cacna1b* (Addgene: 26569) and e37b-Cacna1b (Addgene: 26571) splice isoforms, together with Ca_V_β3 (Addgene: 26574), Ca_V_α_2_δ-1 (Addgene, 26575) and eGFP (Clontech. CA, USA) in tsA201 cells using Lipofectamine 2000 (Invitrogen, CA, USA) as described {Raingo et al., 2007, #119323}.

### 5.3 Electrophysiology

Standard whole-cell voltage-clamp recordings in tsA201 cells were performed as described previously {Andrade et al., 2010, #58833; Raingo et al., 2007, #6309}. The external solution for all recordings contained (in mM) 1 CaCl_2_, 4 MgCl_2_, 10 HEPES, 135 TEA-Cl, pH adjusted to 7.2 with CsOH). Internal solution contained (in mM): 126 CsCl, 10 Cs-EGTA, 1 Cs-EDTA, 10 HEPES, 4 Mg-ATP, pH 7.2 with CsOH. We applied toxin-containing solutions via a microperfusion system using fine glass pipetes placed in close proximity to the cell in order to reduce dead time to within 1 s, the flow rate was controlled by computer-driven solenoids (Warner instruments, VC-66CS) and was set at 0.8 mL/min by adjusting the reservoir height (30 cm). Recording electrode resistence was 2–4 MΩ when filled with internal solution and were coated with Sylgard (Dow Corning) to reduce pipette capacitance. Series resistances (<6 MΩ) were compensated 70–80% with a 10-ms lag time. Calcium currents were evoked by voltage steps from a holding potential of -100 mV, and leak-subtracted online using a P/-4 protocol. We used pClamp version 8.1 software and the Axopatch 200A (Molecular Devices, CA, USA) for data acquisition; data were filtered at 2 kHz and sampled at 20 kHz (–3 dB). All recordings were obtained at room temperatures (22-25 °C). tsA201 cells were typically held at –100 mV to remove closed-state inactivation, and short duration test potentials (20 ms) were applied every 5 s.

### 5.4 Statistical analysis

First, we determined if data follow a normal distribution using the Shapiro-Wilk normality test, data with p > 0.05 were considered normally distributed. Comparisons between groups where all data sets show normal distribution was performed using unpaired two-tailed Student’s *t*-test. Comparison between groups with at least one non normal distribution was performed using Wilcoxon Ran Sum test. For multiple comparisons between varialbles with repeated observations, we used ANOVA repeated measures. We used the R software for all our analysis. Inclusion criteria in our datasets, only successfully recorded cells with steady-state block were included in our analysis and cells where the flow rate was disrupted by ocassional bubles were discarded (1/20). For the kinetic analysis, 2 cells were excluded from the Ca_V_2.2-e37a group because it was not possible to fit one exponential.

## Abbreviations

PnTx3-4: Phoneutria nigriventer toxin PnTx3-4
Phα1β: Phoneutria nigriventer toxin Phα1β
Ca_V_: voltage-gated calcium ion channel
ω-CTX GVIA: ω-conotoxin GVIA
Trpv1: transient receptor potential vanilloid 1

## Author contributions

Author contributuions: Planned experiments (AA, CC-J, DL), performed experiments (AA, CC-J) analyzed the data (AA, CC-J, DL), wrote the manuscript (AA, CC-J, DL), and provided toxins used in these experiments (MVG, MNC, and MHB).

## Acknowledgements

We would like to thank Sylvia Denome for all technical assistance. This work was supported by NIH grants MH099405 (AA) and NS055251 (DL) and CNPq Universal 405175/2018-3 (CCJ, MVG, MNC, MHB) and FAPEMIG Universal APQ-03767-16 (CCJ, MVG, MNC, MHB).

